# RNASeq analysis of a Pax3-expressing myoblast clone *in-vitro* and effect of culture surface stiffness on differentiation

**DOI:** 10.1101/2021.09.05.459022

**Authors:** Louise Richardson, Dapeng Wang, Ruth Hughes, Colin A Johnson, Michelle Peckham

## Abstract

Skeletal muscle satellite cells cultured on soft surfaces (12kPa) show improved differentiation than cells cultured on stiff surfaces (approximately 100kPa). To better understand the reasons for this, we performed an RNA-Seq analysis for a single satellite cell clone (C1F) derived from the H2k^b^-tsA58 immortomouse, which differentiates into myotubes under tightly regulated conditions (withdrawal of γ-interferon, 37°C). As expected, the largest change in overall gene expression occurred at day 1, as cells switch from proliferation to differentiation. Surprisingly, further analysis showed that proliferating C1F cells express Pax3 and not Pax7, confirmed by immunostaining, yet their subsequent differentiation into myotubes is normal, and enhanced on softer surfaces, as evidenced by significantly higher expression levels of myogenic regulatory factors, sarcomeric genes, enhanced fusion and improved myofibrillogenesis. Levels of RNA encoding extracellular matrix structural constituents and related genes were consistently upregulated on hard surfaces, suggesting that a consequence of differentiating satellite cells on hard surfaces is that they attempt to manipulate their niche prior to differentiating. This comprehensive RNASeq dataset will be a useful resource for understanding Pax3 expressing cells.

## Introduction

Satellite cells are the stem cells of skeletal muscle. They lie underneath the basal lamina of the muscle fibre, and transition from a quiescent to an activated state to repair, regenerate or grow muscle fibres in vivo (reviewed in ^1,2^). Following activation, they proliferate to form new daughter satellite cells that replenish the stem cell niche or differentiate into muscle fibres. Satellite cells can be isolated from skeletal muscle and the resulting myoblasts recapitulate the process of muscle differentiation in culture into multinucleated myotubes (reviewed in ^3^).

Most studies of cultured myoblasts use the C2C12 muscle cell line, a subclone of C2 cells originally isolated as a spontaneously transformed myoblast cell line ^4,5^. Differentiation of C2C12 cells is typically initiated by allowing the cells to become confluent and changing the medium from growth to differentiation medium. However, these cells have some differences from primary satellite cells ^3^ including the ability to induce tumour formation if introduced into muscle in vivo ^6^.

An alternative to C2C12 cells, is to use a myoblast clone isolated from the H2k^b^-tsA58 immortomouse ^7^. These cells contain the temperature sensitive mutant of the SV40 large T-antigen (tsA58) under the control of an inducible promoter (*H-2k*^*b*^). The addition of interferon-γ (IFN-γ) drives transcription of the tsA58 gene, and at a temperature of 33°C the T-antigen is stable and promotes cell proliferation. Removing IFN-γ stops transcription and increasing the temperature to 37-39°C leads to degradation of any remaining expressed T-antigen, allowing the myoblasts to exit the cell cycle and differentiate ^8^. These cells have proved valuable in the area of myoblast research as they more faithfully recapitulate the behaviour of primary myoblasts, and do not form tumours when transplanted into mice ^8^. Moreover, these cells can be derived as single clones, with varying behaviour. For example, one clone (2B4) exhibits stem cell behaviour ^9^, while other clones show differences in adult skeletal myosin isoform expression in differentiating myotubes ^10^.

The aim of this research was to perform an extensive differential gene expression analysis of a single myoblast clone isolated from the H2k^b^-tsA58 mouse during differentiation, to characterize changes in muscle gene expression. In addition, we tested how gene expression was affected by culturing these cells on soft surfaces, in which the stiffness matches that of the muscle fibre (approximately 12kPa) compared to hard (approximately 100kPa) glass or plastic surfaces. C2C12 cells differentiate optimally on these soft surfaces ^11^. Finally, the differential gene analysis revealed that the specific myoblast clone we analysed here (C1F) predominantly expresses Pax3 and not Pax7. Although both Pax3 and Pax7 are important for satellite cell formation, Pax3 positive satellite cells are rarer than Pax7 positive satellite cells (reviewed in ^1^), and this has enabled us to explore the differentiation of this rarer type of satellite cell *in vitro*.

## Results

### Overall gene expression is affected by the culture surface

The PCA analysis (Fig 1A) shows that at each day of sampling, the RNA-Seq data for samples from cells cultured on hard surfaces clusters separately to samples from cultures cultured on soft surfaces. This indicates that gene expression patterns within each set of samples are highly correlated and is consistent with the idea that gene expression is affected by the surface on which the cells were grown. Repeats for the same conditions do not completely overlap because of batch-to-batch variation. A hierarchical analysis (Fig 1B) also shows that samples within each treatment (hard vs soft) cluster together but samples from different treatments are less closely linked. Overall, there is a marked difference in gene expression between cells grown on standard hard plastic surfaces compared to soft (approx. 12kPa stiffness) PDMS surfaces.

**Figure 1.**
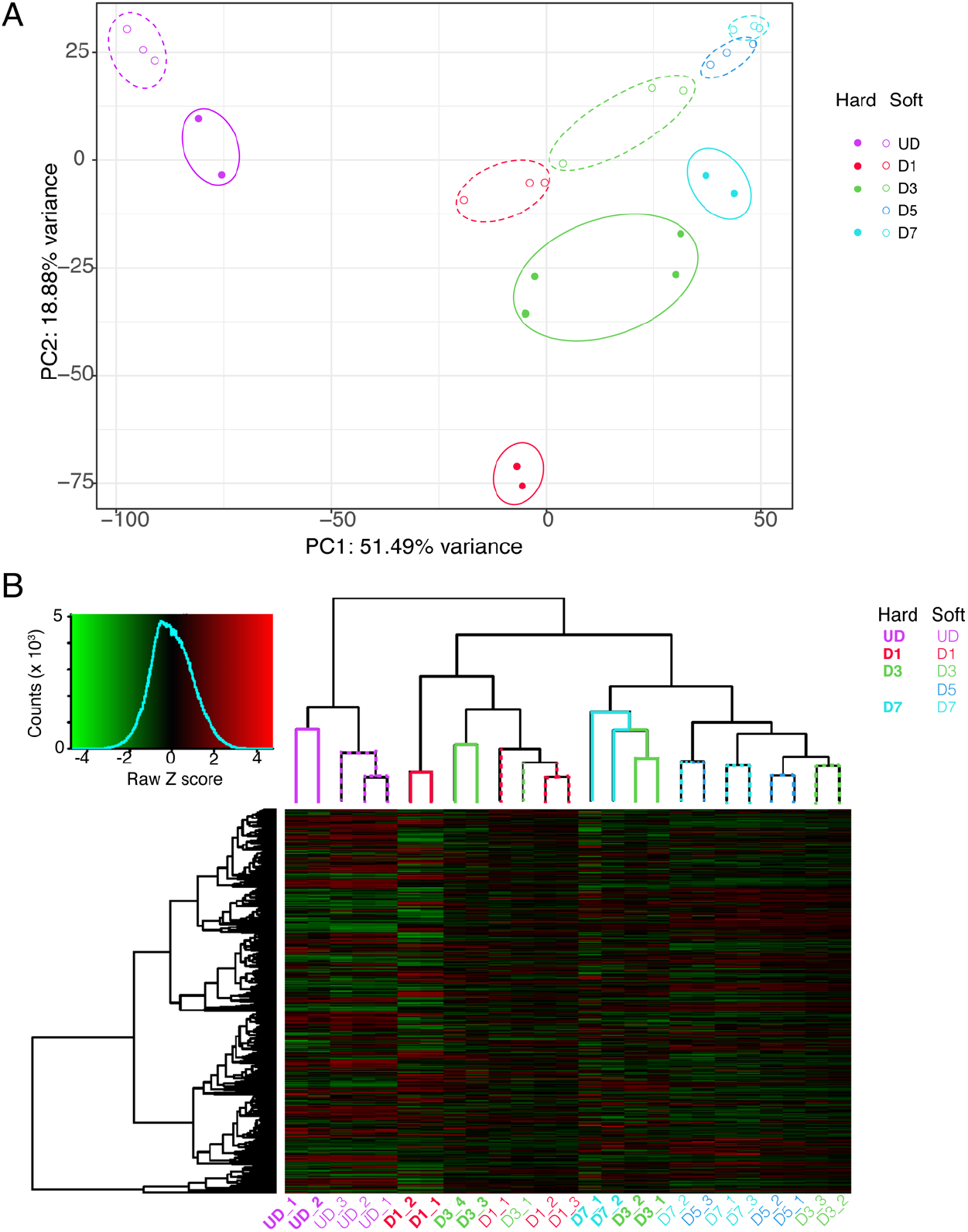
PCA plot and dendrogram for cells differentiating on hard or soft surfaces. **(A)** shows the PCA analysis for each set of samples at each time point. ‘Hard’ represents hard surfaces (approximately 100kPa stiffness). ‘Soft’ represents soft surfaces (approximately 12kPa stiffness). At each time point, the values for samples on hard surfaces and soft surfaces are shown. **(B)** Hierarchical clustering of gene expression for cells grown on hard or soft surfaces at different each time point. The branch length at the top of the figures indicates the level of dissimilarity between sample clusters (those with shorter branches have a higher degree of similarity). Each individual branch represents a group of related genes expressed within the corresponding sample labelled underneath

### Comparison of overall gene expression between hard and soft surfaces over time

We next performed an analysis of the overall variation in gene expression levels between cells grown on soft and hard surfaces at each time point (Fig. 2A-D). A global analysis of the change in expression levels across genes that show a significant change (_padj_<0.05; log_2_FoldChange <-1 or >1), showed that the largest difference in gene expression occurred at day 1 (D1) of differentiation. This is consistent with the switch to terminal differentiation into myotubes at D1. DEGpatterns analysis (see methods) of the RNASeq data on hard soft surfaces showed a higher number of clusters for changes to gene expression in cells cultured on soft surfaces compared to those cultured on hard surfaces (Supplemental Fig.S1).

**Figure 2.**
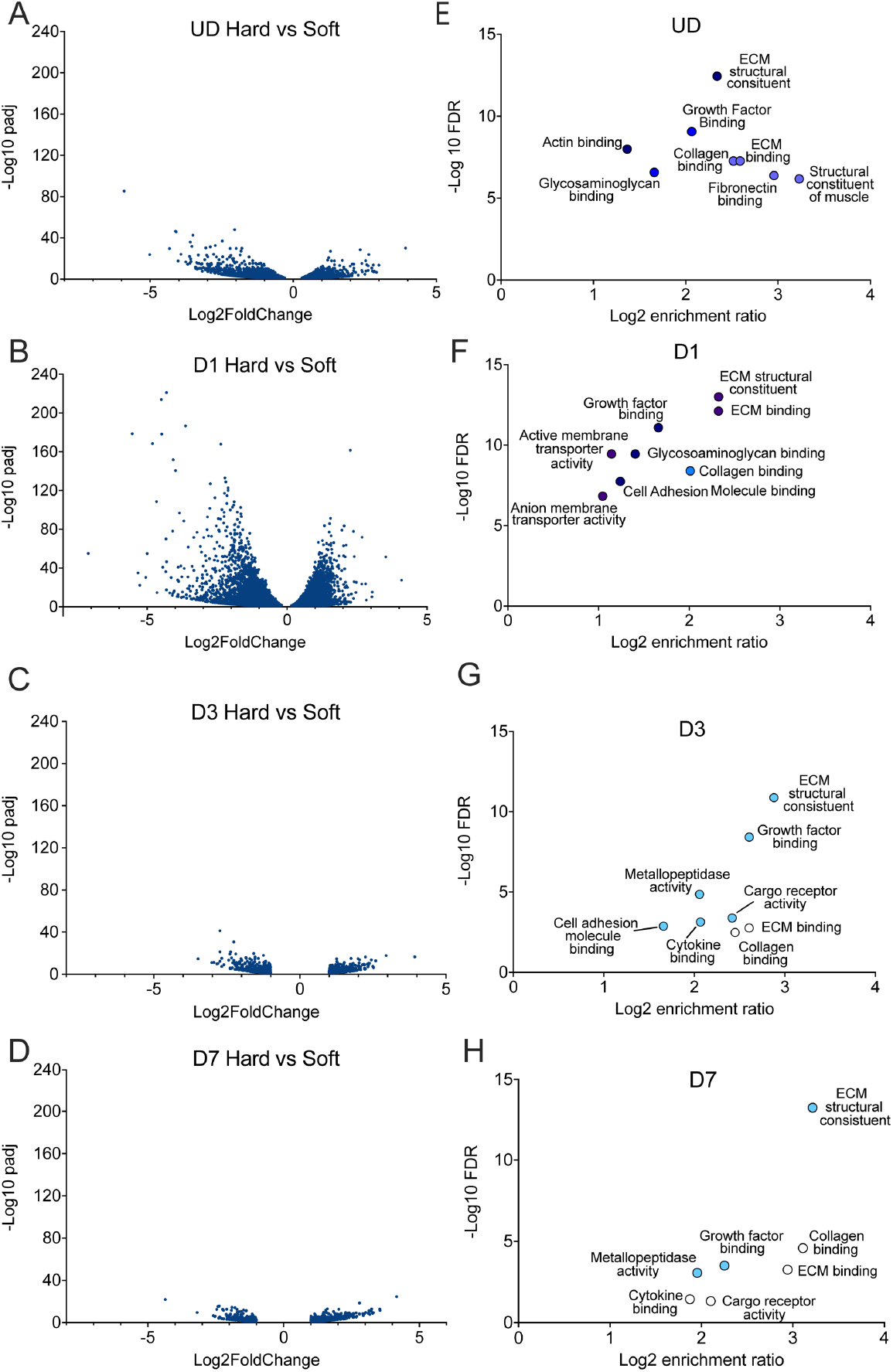
Global comparison of changes in gene expression levels at each time point between hard and soft surfaces. A-D show volcano plots for gene expression where the log_2_fold change is either >1 or <-1, and where the padj is <0.05. E-H show the results from a WebGestalt analysis for enriched gene sets (molecular function) at each time point, for a comparison of hard with soft surfaces, down-regulated genes (e.g lower expression on soft surfaces). The log10 false discovery rates (FDR) are shown for those with a p value of <0.05. The symbol colour for each gene set is approximately related to the number of enriched genes within each gene set from dark blue (>100) to white (<20).

A WebGestalt analysis of genes expressed at > 2 fold higher (log_2_FoldChange>1) levels on hard than on soft surfaces (p_adj_ <0.05) revealed that RNA expression for genes associated with the structural constituent of muscle and those for extracellular matrix (ECM) were higher on hard surfaces in undifferentiated cells (Fig. 2E). Genes encoding ECM structural constituents, ECM binding and growth factor binding were enriched for cells cultured on hard surfaces across the time course (Fig. 2E-H). To explore these changes further we first analysed expression of transcription factor genes associated with myogenesis and genes associated with myofibrillogenesis (formation of muscle sarcomeres).

### Changes in myogenic transcription factor expression between hard and soft surfaces

Quiescent and newly activated satellite cells typically express one or both of two paired box transcription factor genes, Pax3 and Pax7. Pax7 is more commonly expressed in satellite cells than Pax3 ^1^ and muscle cell lines such as C2C12 express Pax7 ^12^. The RNA-Seq data demonstrated that undifferentiated (UD) C1F cells on soft and hard surfaces express Pax3 (Fig. 3A) and that levels of Pax3 decrease as the cells differentiate, as expected (Fig. 2A). A differential gene analysis that independently compared levels of Pax3 at each day (from D1 to D7) to those in undifferentiated cells (UD, at Day 0) showed that levels of Pax3 were significantly lower at each day (D1-D7) of differentiation compared to UD cells (Fig. 3A, B) on soft surfaces. In contrast, Pax3 levels did not significantly decrease on hard surfaces compared to UD until D7. Directly comparing Pax3 expression levels between cells on soft and hard surfaces at the same time point (Day 1) showed a small but significant increase in expression of Pax3 on soft surfaces compared to hard surfaces (Fig. 3C). This suggests that soft surfaces might promote higher levels of Pax3 expression.

**Figure 3:**
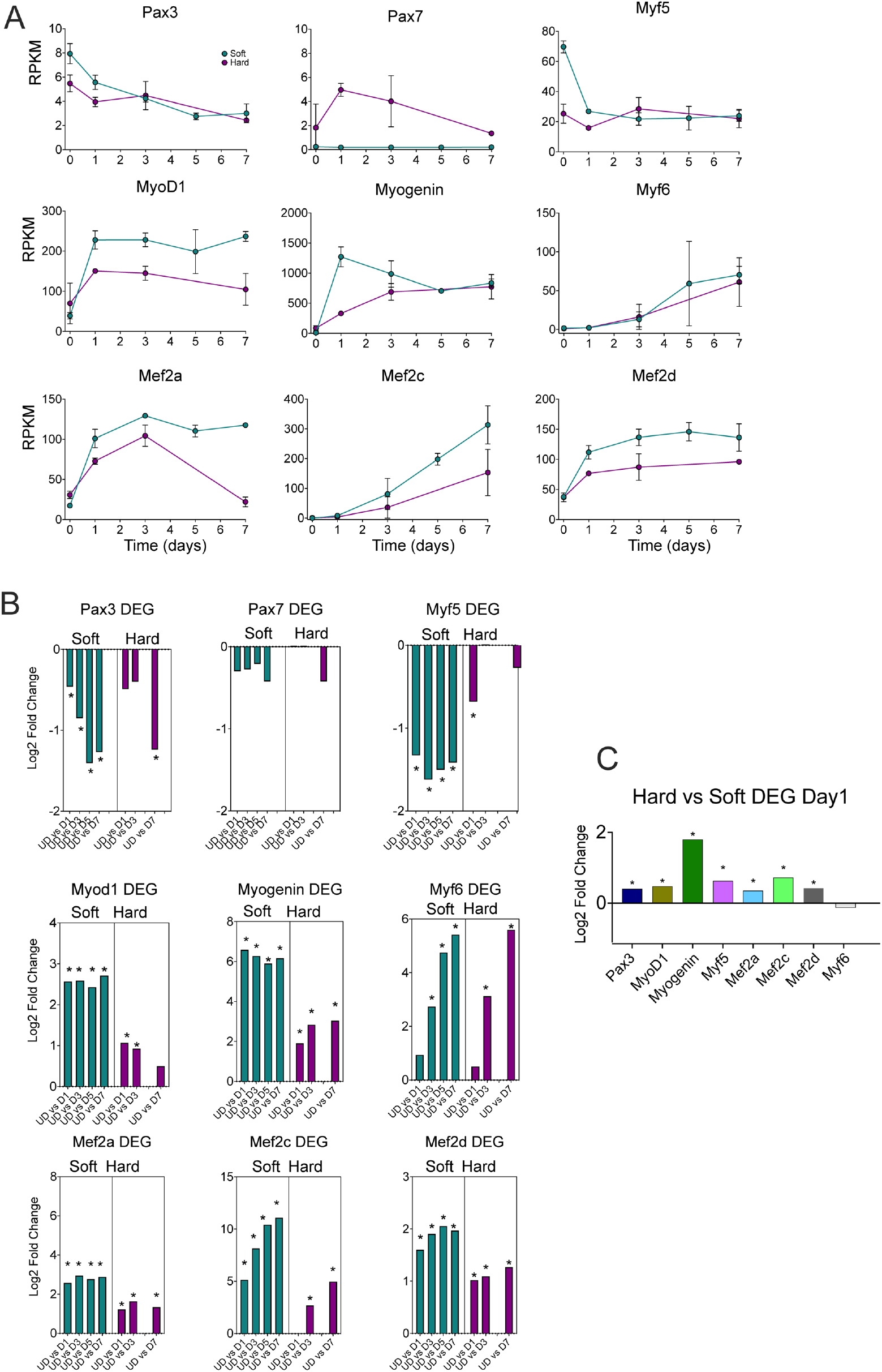
Expression of myogenic factors on soft and hard surfaces in C1F cells. **A:** shows the RPKM values for each time point samples for soft (green) and hard (magenta) surfaces, from undifferentiated cells (UD) to Day 7 of differentiation (D7). Error bars show the standard deviation (S.D.). **B**: shows a differential gene analysis (DEG) for each of the myogenic factors over time, comparing UD with D1 etc, for soft (green) and hard (magenta) surfaces, * indicates p_adj_ <0.05. C shows the results for a DEG comparing expression for each of the main myogenic factors for cells cultures on hard versus soft surfaces at day 1, when the main changes in myogenic factors are observed. * Indicates p_adj_<0.05.

In contrast, levels of Pax7 in cells cultured on soft surfaces were very low compared to cells on hard surfaces (Fig. 3A). Expression of Pax7 appears to be higher for cells grown on hard surfaces and appears to increase from UD to D1 and D3 (Fig. 3A,B). However, RPKM (reads per kilobase per million mapped reads) values for Pax7 are 50% lower than for Pax3 even on hard surfaces (Fig. 3A), a differential gene analysis showed no significant change in expression between UD cells and D1-7 cells (Fig. 3B) and it would be unusual for Pax7 to be expressed at higher levels at D1 and D3 compared to undifferentiating cells as appears to be the case for cells on hard surfaces. A direct comparison of Pax7 expression between soft and hard surfaces at day 1 (Fig. 3C) does show a significantly higher expression level for Pax7 in cells cultured on hard surfaces. Overall, these results suggest that the C1F cell line is unusual in that it predominantly expresses Pax3 rather than Pax7.

Next, we investigated the expression patterns of the four myogenic regulatory factors Myf5, MyoD, myogenin and MRF4 (Myf6), which belong to the helix-loop-helix family of transcription factors ^13,14^. The expression of Myf5, MyoD, myogenin and Myf6 is hierarchical, with Myf5 and MyoD being expressed first, followed by myogenin and Mrf4 (Myf6) ^13^. Similar to Pax7 and Pax3, Myf5 can also be expressed in quiescent satellite cells and/or throughout differentiation ^15^.

The expression levels of Myf5 decreased significantly in differentiating cells at D1 to D7 compared to UD cells for cells grown on both soft and hard surfaces (Fig.3 A,B) showing a similar trend in expression to Pax3. The expression levels of MyoD increased at D1 and then remained roughly similar on both soft and hard surfaces (Fig. 3A). The differential gene analysis showed that the expression levels of MyoD were increased significantly at D1-D3 compared to UD cells on both soft and hard surfaces and were also increased significantly at D5 and D7 compared to UD on soft surfaces. Myogenin expression was increased significantly in cells cultured on both soft and hard surfaces at D1-D7 compared to undifferentiated cells (Fig. 3A,B). However, the increase in myogenin expression was much higher at D1 on soft, compared to hard surfaces (Fig. 3C). Myf6 expression was significantly increased at later time points, from D3 onwards, compared to UD cells (Fig. 3A,B) for cells cultured on both soft and hard surfaces. This later rise is consistent with previous work ^15^.

The Mef2 family of genes (Mef2a,b,c and d) encode myogenic transcription factors that belong to the MADS family of transcription factors ^16^. Mef2c can act synergistically with MyoD and myogenin to promote myogenesis, while Mef2a and d can act synergistically with MyoD and this is mediated through direct interaction of the two proteins ^17^. Mef2c has also been shown to be important for the maintenance of sarcomere integrity, by regulating transcription of the sarcomeric gene myomesin ^18^. We found that levels of Mef2a,c and d increased significantly from UD cells to D1-7 cells on both hard and soft surfaces although the magnitude of the log_2_fold change was higher on soft surfaces (Fig. 3A,B).

Directly comparing gene expression for all myogenic transcription factors at day 1 showed expression is significantly higher for cells on soft, compared to hard surfaces, with the exception of Myf6 (Mrf4) (Fig 3C). Strikingly, the increase in expression levels for myogenin between UD and D1 was much higher for cells on soft surfaces compared to those on hard surfaces (Fig. 3C). As myogenin is a master regulator of differentiation, this indicates that the cells on soft surfaces may be differentiating into myotubes at a faster rate than those on hard surfaces.

To confirm changes in RNA expression, we analysed C1F cells fixed and stained for a subset of myogenic transcription factors: Pax3, MyoD and myogenin from undifferentiated cells to day 7 of differentiation (Fig. 4). Levels of Pax7 were undetectable in CIF cells (Fig. 4a). Levels of Pax3 were significantly higher in CIF cells on soft compared to hard surfaces in undifferentiated cells and at D1, and then were undetectable from D3-D7. Levels of MyoD increased at D1 and then gradually declined, with levels significantly higher in cells on soft surfaces from D1-D7. Levels of myogenin increased at D1 in cells on soft surfaces, and then gradually declined, while levels of myogenin in cells on hard surfaces increased later at D3 and levels were significantly higher in cells cultured on soft surfaces compared to hard surfaces at D1 and D3. Overall, these results are in broad agreement with the RNASeq analysis and confirm that the C1F cell express Pax3 and not Pax7.

**Figure 4.**
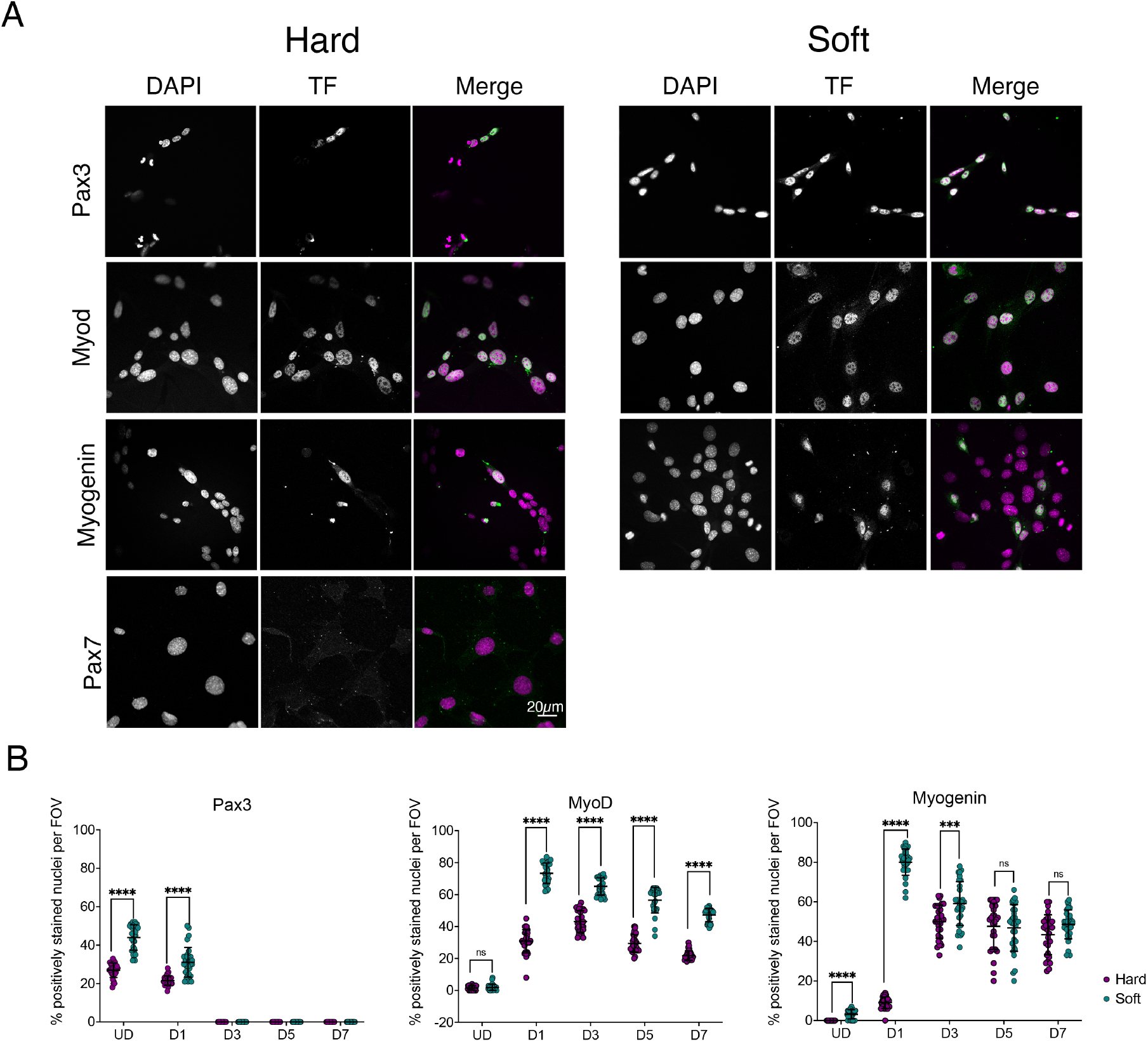
Myogenic transcription factor expression in cells on soft and hard surfaces. A. example images of cells stained for DAPI (magenta) and specific transcription factors (green). B. Quantification of the number of nuclei staining positively for each transcription factor (Pax3, MyoD and myogenin) over time on hard (magenta) and soft (green) surfaces. **** p<0.0001, *** p<0.001. ns: non-significant

The rapid increase in myogenin mRNA for cells on soft surfaces at D1 further suggested that fusion might be promoted on soft surfaces. To investigate this further, we fixed and stained the cells for skeletal myosin heavy chain at different time points, and quantified fusion on the two types of surfaces (Fig. 5). We found that fusion index was indeed increased on soft surfaces (Fig 5B). This is consistent with the early rise in myogenin expression observed for cells cultured on soft surfaces. Imaging the organisation of skeletal muscle myosin heavy chain in D5 myotubes shows improved organisation and better alignment of sarcomeres in cells on soft surfaces compared to hard surfaces (Fig. 5C)

**Figure 5.**
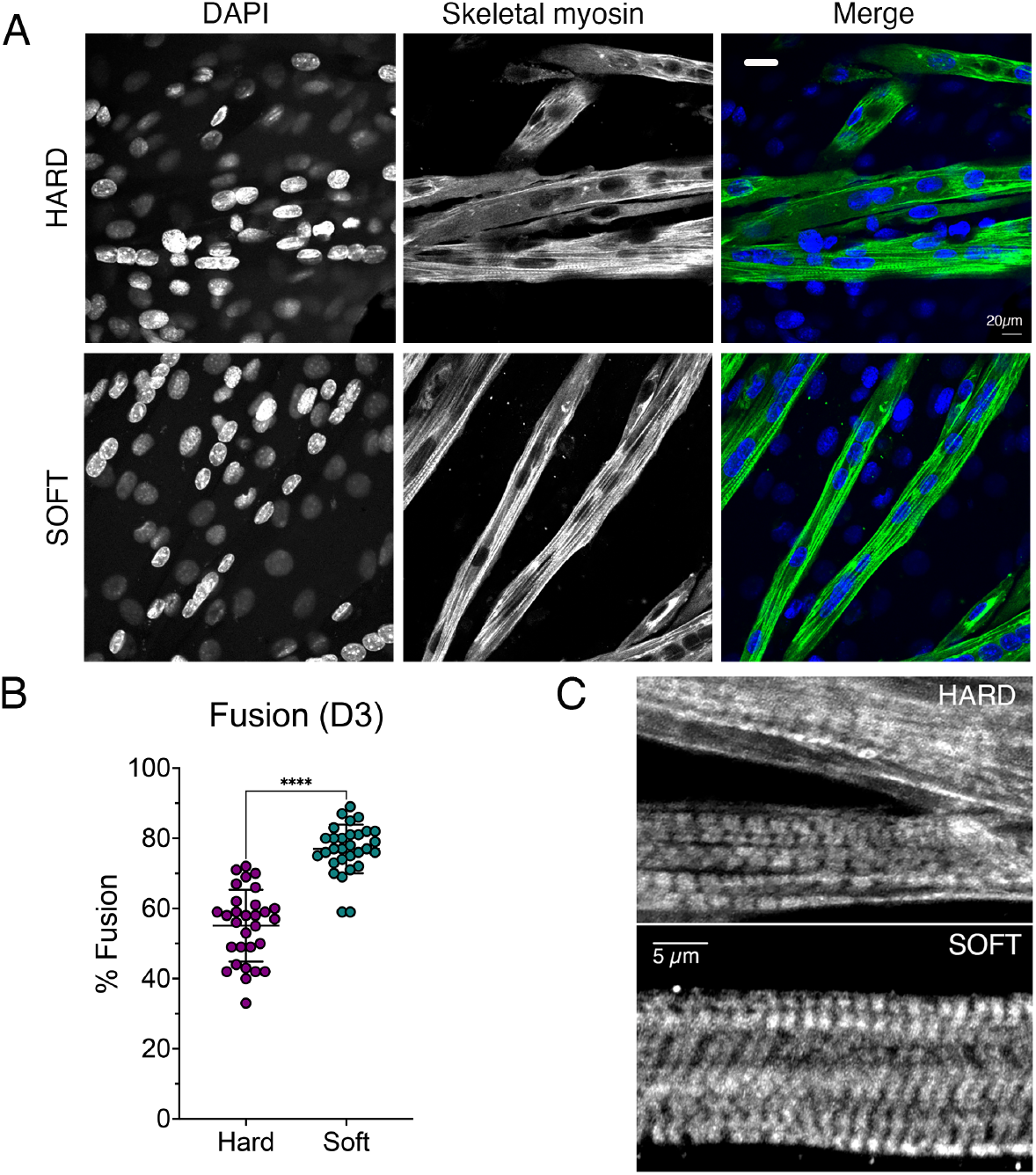
Fusion and differentiation on hard and soft surfaces. A: representative fields of view for differentiating cells showing nuclei (blue) and myotubes stained for skeletal myosin (green). B: fusion index measured for cells on hard (magenta) and soft (green) surfaces at D3. C: Images of myotubes at day 5 on hard and soft surfaces stained for skeletal myosin, to show sarcomeric organisation.

Next, we further analysed the RNASeq data to determine the general trend for expression of a selection of sarcomeric proteins, given the improved myofibrillogenesis on soft surfaces shown here and in agreement with previous work using C2C12 cells ^11^. We focussed on key structural proteins, such as skeletal (Acta1) and cardiac actin (Actc1), actinin-2 (Actn2: found in the Z-disc), myosin heavy chain genes (Myh3 – embryonic, Myh8 – perinatal, and Myh7 – slow/β-cardiac), titin (Ttn), nebulin (Neb), t-cap, myosin binding protein-C (Mybpc) and Unc45b, a specific striated muscle chaperone, important for the folding of skeletal myosin heavy chain ^19^. The RNA expression levels for all of these genes generally increased as the cells differentiated, consistent with the myogenic nature of the C1F clone (Fig. 4). However, changes were more marked for cells cultured on soft surfaces (Fig. 6A,B).

**Figure 6:**
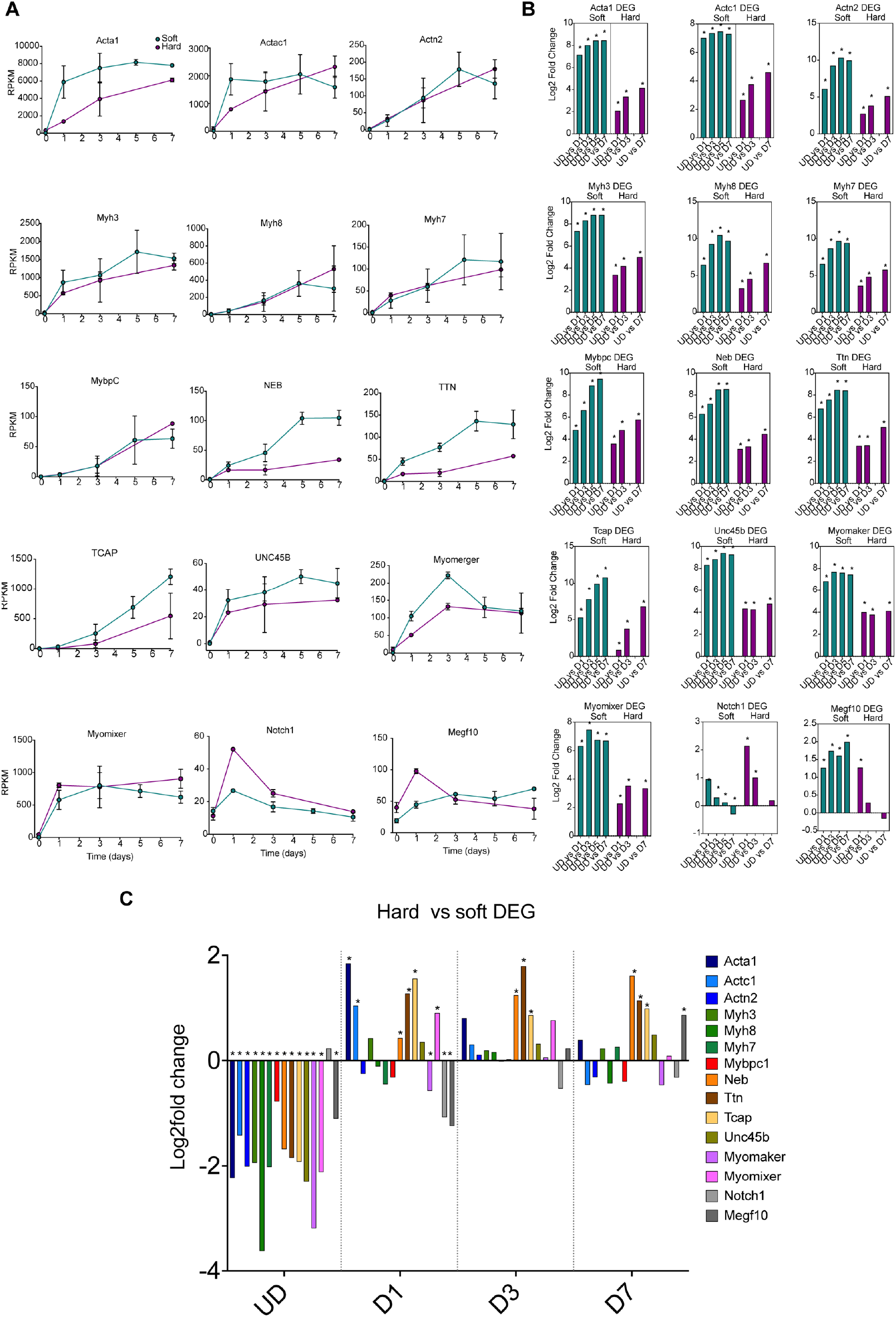
Expression of sarcomeric genes on soft and hard surfaces in C1F cells. A: shows the RPKM values for each time point samples for soft (green) and hard (magenta) surfaces, from undifferentiated cells (UD) to Day 7 of differentiation (D7). Error bars show the standard deviation (S.D.). B shows a differential gene analysis (DEG) for each of the genes over time, comparing UD with D1 etc, for soft (green) and hard (magenta) surfaces, * p_adj_ <0.05. C shows the results for a DEG comparing hard with soft surfaces at each day from UD to D7, * p_adj_ <0.05.

Directly comparing soft with hard surfaces (Fig. 6C) showed that RNA expression levels for many sarcomeric genes are lower on soft than for hard surfaces in undifferentiated cells. This suggests that growth on soft surfaces might tend to inhibit differentiation. This difference in expression quickly reversed on day 1 for many of these genes, with expression levels significantly higher in cells cultured on soft surfaces, concomitant with the increase in myogenin levels (Fig. 3) as cells switch to differentiation.

In addition to the sarcomeric genes, we investigated the expression pattern for two key genes that encode membrane proteins myomixer (Gm7325) and myomerger (Tmem8c). These two genes are important in myoblast fusion, required to form multinucleated muscle myotubes ^20^. The expression of both genes is increased significantly from D1-7 on both soft and hard surfaces, compared to undifferentiated cells (Fig. 6A,B). Expression of myomixer is increased significantly at D1 on soft surfaces compared to hard surfaces (Fig. 6C), potentially helping to explain the increased ability of C1F cells to fuse on soft surfaces.

Notch1 signalling is also known to be important in muscle myogenesis. Notch1 is upregulated when satellite cells are activated and promotes proliferation ^21^. Overexpression of Notch1 can inhibit myogenesis ^22^. We found that expression of Notch1 increased significantly between UD and D1 of differentiation on both hard and soft surfaces (Fig.6A,B). Moreover, there was a significantly higher level of expression of Notch1 in cells cultured on hard surfaces at D1 (Fig. 6C). This is unexpected because Notch1 expression should decrease at D1, as the time cells cease proliferating and start differentiating. MEGF10 has been suggested to interact with Notch1 and to be significantly downregulated when C2C12 cells were induced to differentiate by serum withdrawal ^23^. We found that expression levels of MEGF10 increased significantly between undifferentiated cells and D1 on both soft and hard surfaces. However, on soft surfaces, MEGF10 levels remain significantly elevated at D3-7 compared to UD cells, while on hard surfaces, MEGF10 levels do not. MEGF10 levels are significantly increased for cells cultured on hard surfaces at D1 (Fig. 6C) and then significantly increased for cells cultured on soft surfaces at D7. These expression patterns argue against the role of MEGF10 in promoting proliferation in combination with Notch1.

Overall, this analysis of gene expression demonstrates that Pax3 positive C1F cells express a range of markers for myogenic differentiation, with a pattern consistent with differentiation of myoblasts into multinucleated myotubes.

### Differential gene expression analysis of collagen genes

The WebGestalt analysis showed that, growing cells on hard surfaces increased gene expression for a range of ECM and associated proteins, as well as the receptors that bind these proteins. This raises the possibility that the cells cultured on hard surfaces attempt to form their own niche, through expression of ECM proteins, to enable them to differentiate on hard surfaces, and could help to explain the delay in expression of myogenic genes.

To explore the results from the WebGestalt analysis further, we analysed the expression of collagen (COL) genes found in the ECM structural constituent category between hard and soft surfaces at each time point. This analysis focused on genes with the highest RPKM values and known to be associated with skeletal muscle ECM ^24^. Almost all the COL genes analysed showed a high early peak in RNA expression at D1 in cells cultured and a generally higher overall RNA expression of collagen genes (Fig. 7A) on hard surfaces. A differential gene expression analysis (DE) that compared expression levels at each day of differentiation with that in undifferentiated cells grown on the same type of surface showed similar trends for collagen expression for most collagen genes (both Col1 genes, Col3a1, Col5a1 and 2, both Col8 genes, Col12a1, Col16a1 and Col18a1: Fig. 7B). A differential expression analysis comparing expression at each time point between soft and hard surfaces, showed higher levels of collagen in cells cultured on hard surfaces (Fig. 7C).

**Figure 7:**
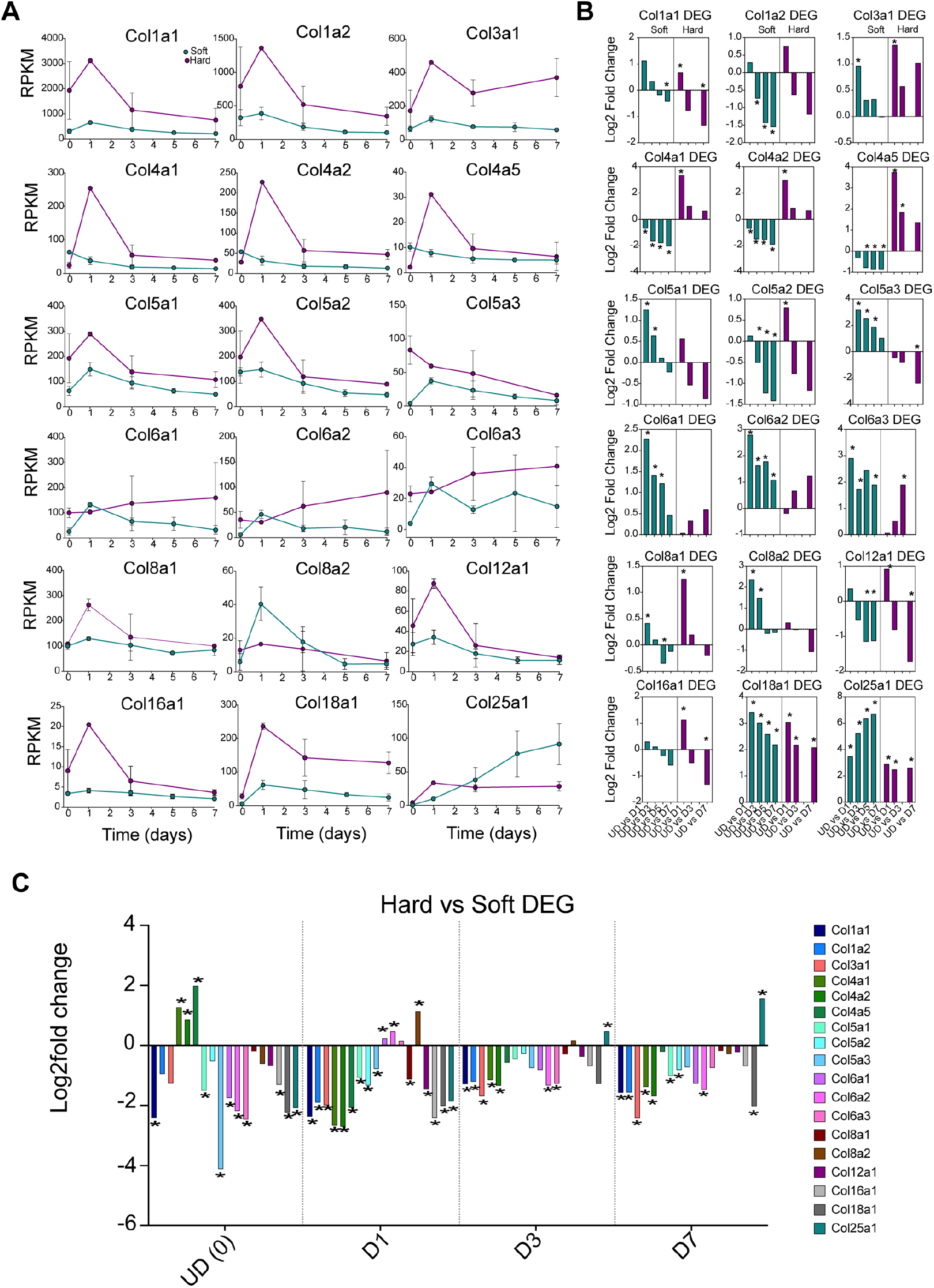
Comparison of gene expression for collagens important in muscle on hard and soft surfaces over time. A: RPKM plots for hard (magenta) and soft (green) surfaces for each of the collagens, over time. Day 0 (D0) represents undifferentiated cells (UD). Days 1-7 represent the time course of differentiation. B: Differential gene analysis for each day of differentiation (D1-7) to undifferentiated cells (UD) for cells cultured on hard and on soft surfaces, *indicates significant change in Log_2_fold expression (p_adj_ value <0.05). C: differential gene analysis for expression levels of each collagen isoform between hard and soft surfaces, at each day (from UD to D7), * indicates significant change in Log_2_fold expression (p_adj_ value <0.05)

While the general trend in differential expression of collagen genes was similar, some differences were observed. The RNA expression profiles of COL4 genes (Collagen IV: Col4A1, 4A2 and 4A5) between cells cultured on soft and hard surfaces. On hard surfaces, the RNA expression levels for Col4 peaked at day 1, and levels were higher than that found for cells cultured on soft surfaces. A differential gene analysis showed RNA levels from D1-D7 were lower than in UD cells for cells cultured on soft surfaces (Fig. 7B). In contrast RNA levels increased at D1 for cells cultured on hard surfaces before then decreasing from D3-D7, but levels remained higher than in UD cells. A direct comparison between soft and hard surfaces showed that Col4 RNA levels for each of these three isoforms were higher in undifferentiated cells on soft surfaces compared to those on hard surfaces (Fig. 7C). Also of note is that there is an expression peak of Col8a2 on soft surfaces, but not on hard surfaces, the reverse of the general trend. Finally, Col25a1 continues to increase from UD to D7 on soft surfaces but after increasing at D1, it stays about the same on hard surfaces.

Collagen IV is part of the basement membrane, a thin sheath of connective material that surrounds skeletal muscle, and other cell types ^25^. A heterotrimer composed of two α1 (Col4a) and one α2 molecules is found in many types of basement membranes, including that of muscle ^26^. The increased expression of Col4 genes in cells cultured on hard surfaces at day 1 of differentiation suggests that the hard surface is promoting basal lamina formation more strongly than the soft surface, possibly to counteract the hardness of the culture surface. The reasons for differences in expression profiles for Col8a1 and Col25a1 are unclear.

### Differential gene expression analysis of laminin, nidogen and integrin genes

We performed a similar differential gene expression analysis for laminin and nidogen, two proteins also found in basal lamina alongside Collagen IV, and integrins, membrane proteins that bind ECM proteins. Laminins (Lama2, Lama5, Lamb1, Lamb2) also show a clear peak in expression at day 1 on hard surfaces (Fig. 8A) consistent with the idea that the cells on hard surfaces are stimulated more strongly to create their own ECM niche. Nidogen (Nid1) also showed a higher peak of expression at day 1 on harder surfaces (Fig. 8A). Of the integrin genes with the highest levels of RNA (as judged by RPKM levels) integrins α5 (Itga5), α7 (Itga7) and β1 (Itgb1) had a peak of expression at day 1 on hard surfaces, which was higher than that on soft surfaces (Fig. 8A,C). Integrin α5, β1, binds to fibronectin (FN1), and its levels decrease as myoblast fusion and differentiation progress, as reported previously ^27^. Integrin α7, β1 is the main laminin receptor on myoblasts and myotubes and has been shown to be important for myoblast fusion ^28^. A differential gene analysis over time, on hard and on soft surfaces showed a general decrease in expression for the majority of these genes.

**Figure 8:**
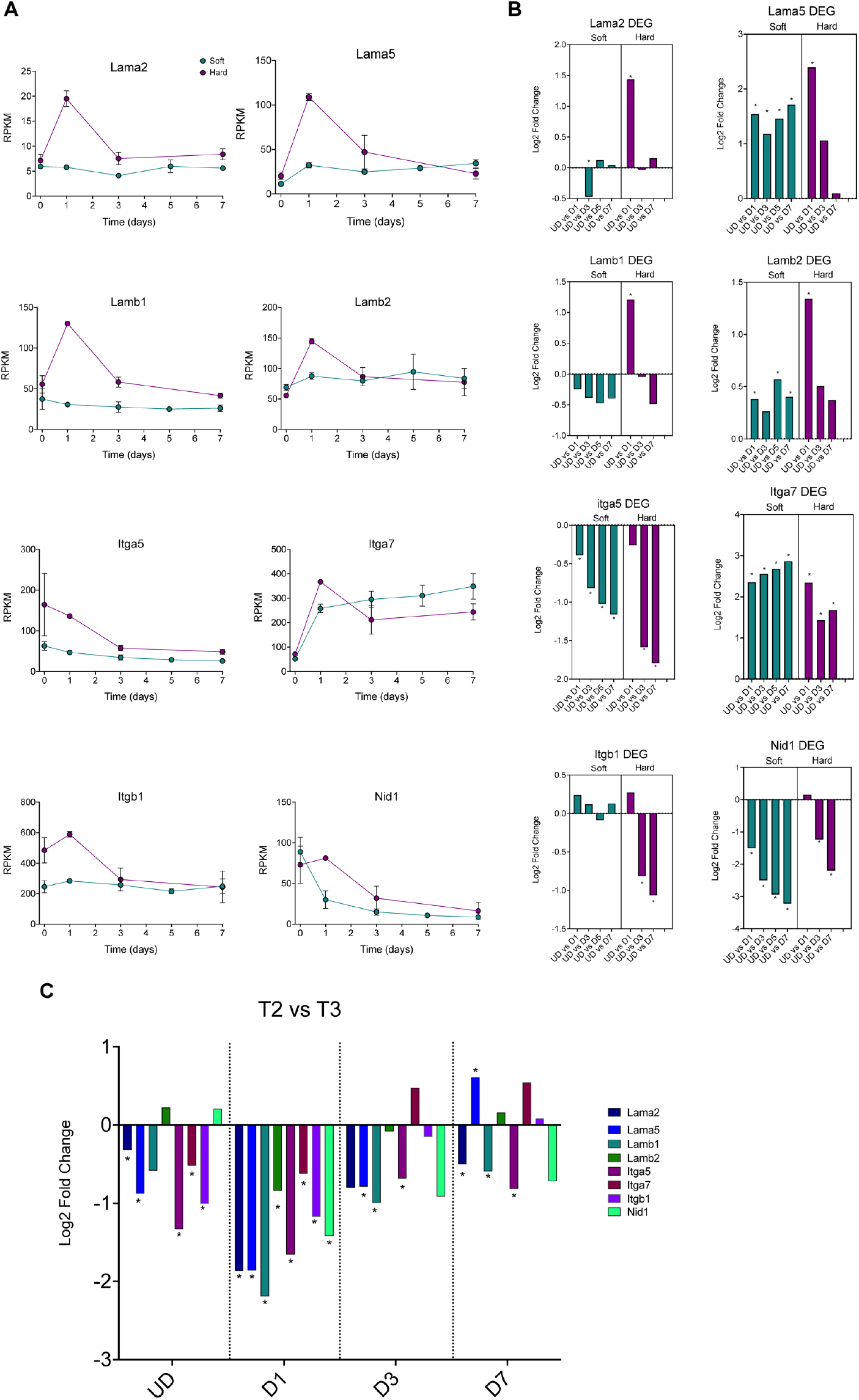
Comparison of gene expression for laminins, integrins and Nid1 (nidogen) on hard and soft surfaces over time. A: RPKM plots for hard (magenta) and soft (green) surfaces for each of the genes of interest, over time. Day 0 (D0) represents undifferentiated cells (UD). Days 1-7 represent the time course of differentiation. B: Differential gene analysis comparing each day of differentiation (D1-7) to undifferentiated cells (UD) for cells cultured on hard and on soft surfaces, *indicates significant change in log_2_fold expression (p_adj_ value <0.05). C: differential gene analysis for expression levels of each collagen isoform between hard and soft surfaces, at each day (from UD to D7), * indicates significant change in log_2_fold expression (p_adj_ value <0.05)

## Discussion

These data show that a clonal cell line of myoblasts, C1F, derived from the H2k^b^-tsA58 immortomouse predominantly expresses *Pax3* and not *Pax7* and yet follows the same differentiation pathway observed for *Pax7* expressing cells ^15^. Our findings thus support the evidence that despite the differences in roles of these two genes, *Pax3* and *Pax7* are capable of triggering a myogenic programme that follows the same transcriptional pattern leading to muscle maturation ^29^.

The C1F cells differentiate better on softer (∼12kPa) surfaces than on hard surfaces (plastic or glass), consistent with previous observations for the C2C12 cell line ^30^. The key difference between differentiation on soft and hard surfaces appears to be an earlier higher expression of two myogenic regulatory factors, MyoD and myogenin, which indicate that cells begin to differentiate earlier on softer surfaces. The increase in MyoD agrees with previous reports of a precocious rise in *MyoD* on soft surfaces, alongside overall greater expression by cells ^31^. Since *MyoD* activation is necessary for the specification of terminal differentiation and triggers *myogenin* activation, the entire myogenic process is expedited. Not surprisingly then, we also find an earlier peak in the rise of myogenin on softer surfaces. Our results are also consistent with a previous RNA-Seq analysis for C2C12 cells that showed a greater increase in expression of genes related to sarcomere formation and muscle maturation on micropatterned soft surfaces than non-micropatterned, compared to hard plastic^31^. Interestingly, expression of sarcomeric genes appeared to be supressed in proliferating cells on soft surfaces, compared to those on hard surfaces. This may be consistent with the finding that substrate stiffness can affect satellite cell self-renewal in vitro^32^.

The PCA analysis showed that RNA expression for cells grown on soft surfaces were clustered separately to cells grown on hard surfaces at each time point. A volcano plot for upregulated and downregulated genes at each time point (UD-D7) demonstrated that the change in gene expression was greatest at D1, where the cells are switching from proliferation to differentiation. A WebGestalt analysis of genes upregulated in cell culture on hard surfaces, compared to soft surfaces, suggested that genes coding for extracellular matrix proteins were highly enriched in cells cultured on hard surfaces. This suggests that cells cultured on hard surfaces respond by upregulating expression of ECM genes, possibly in an attempt to manipulate their ECM to facilitate differentiation, which then occurs, but at a slower rate than on soft surfaces.

Overall, the RNA-Seq data here should be a valuable resource for interrogating changes to expression of differentiation Pax3 positive cells, and the effects of culture on soft and hard surfaces on gene expression.

## Materials and Methods

### Preparation of PDMS surfaces

Synthetic culture surfaces were fabricated using Sylgard 184 (Dow Corning, US) elastomer and curing agent. When combined, the elastomer and curing agent form a silicone-based organic polymer known as polydimethylsiloxane (PDMS). A curing agent to elastomer ratio of 1:50 was used to achieve an elastic modulus of about 12kPa (as reported in supplemental information in ref ^33^). Using a w/w ratio, components were weighed into separate glass beakers and autoclaved. Elastomer and curing agent were then mixed together thoroughly with a sterile metal spatula before transfer to a 15ml falcon tube. The PDMS mixture was centrifuged at 1000 x g for 2 mins to degas, and a snipped pipette tip was then used to dispense 1ml of the mixture onto the surface of a 60mm diameter Nunc™ petri dish. The PDMS was carefully smoothed across the entirety of the surface with a sterile metal spatula and left to set at room temperature for 48 hours.

To improve the adherence, fusion and differentiation of cells cultured on the PDMS surfaces, collagen was crosslinked onto the surface ^34^. Collagen solution was prepared by mixing 0.1% [w/v] solution of type 1 collagen from calf skin (Sigma) diluted 1 in 10 with sterile water. Sulfo-SANPAH (N-sulphosuccinimidyl-6-(4′-azido-2′-nitrophenylamino) hexanoate), a hetero-bifunctional crosslinking agent was prepared as a 0.5mg/ml solution with 50mM HEPES buffer at pH 8.5. 3ml Sulfo-SANPAH solution was added to each coated petri dish, to cover the surface. Dishes were then irradiated with a UV source for 8 mins to activate the crosslinker. After exposure, spent Sulfo-SANPAH was aspirated and wells washed with HEPES buffered solution. These steps were then repeated a second time. Next, 3ml collagen solution was then applied to each surface and allowed to adhere for 4hrs at room temperature, before being washed 3 times with PBS. As a comparison to the soft surfaces, non-PDMS coated standard petri dishes were used, coated with the collagen solution for 4 hours at room temperature.

### Growth, proliferation and Isolation of RNA from cultured cells

To determine levels of gene expression on standard versus PDMS surfaces, RNA was isolated from C1F cells at different stages of differentiation. C1F myoblasts were originally isolated from the hindlimb muscles of neonatal (1 day old) H2k^b^-tsA58 immortomice ^8^ and shown to differentiate into myotubes ^10^. C1F myoblasts, at passage 3, were recovered from storage, and proliferated in medium composed of 1 x DMEM with GlutaMAX supplemented with 20% heat inactivated foetal bovine serum (FBS, Gibco), 2% chick embryo extract (CEE,E.G.G. Technologies) and 1% (v/v) Penicillin/Streptomycin (P/S, Gibco), and 20 U/ml gamma interferon (γFN, Life Technologies) ^8^, at at 33°C with 10% CO_2_. For the RNA-Seq experiments, cells were seeded onto standard or PDMS coated 60mm Nunc™ petri dishes at a density of 2×10^5^ cells/ml and incubated for 24 hours before harvesting (UD). To differentiate the cells, the growth medium was exchanged for differentiation medium, composed of 1X DMEM + GlutaMAX (Gibco) supplemented with 4% horse serum (Gibco), 1% CEE and 1% (v/v) P/S, and lacking γFN, the incubation temperature was raised to 37°C, and CO_2_ levels reduced to 5%. Cells were harvested at days 1, 3, and 7 of differentiation. Cells on the softer surfaces were additionally harvested at day 5 of differentiation. To harvest the cells, the medium was removed and cells were lysed directly with TRIzol™ (Invitrogen) reagent, by adding 1ml TRIzol™ to each dish, and using a cell scraper to remove the adherent cells into solution. The resulting cell lysate was mixed thoroughly by pipette, placed into labelled Eppendorf tubes and mixed further by vortex. The lysed cell suspensions were stored at -80°C until they were sent for processing.

### Sample processing

Processing of C1F samples was performed by the Translational Genomics Unit, St James’s University Hospital, using their standard genomics workflow. Briefly, RNA samples were treated with a TURBO DNA-free™ Kit (Ambion Inc.) using conditions recommended by the manufacturers, and then cleaned with an RNA Clean & Concentrator™-5 spin column (Zymo Research Corp.) RNA was tested for quality and yield using a NanoDrop 1000 spectrophotometer and an Agilent 2100 Bioanalyzer.

To minimize bubble PCR artefacts, 100 ng of purified total RNA was used in library preparation, following the TruSeq Illumina protocol. In brief, RNA was polyA-selected, chemically fragmented to about 200 nt in size, and cDNA synthesized using random hexamer primers. Each individual library received a unique Illumina barcode. RNA-Seq was performed on an Illumina HiSeq 3000 instrument using 151 bp paired-end reads. The FASTQ files for the RNA-Seq samples are available from Sequence Read Archive (SRA) under accession number PRJNA682314.

The quality of paired-end FASTQ files was checked through FastQC (https://www.bioinformatics.babraham.ac.uk/projects/fastqc/). Cutadapt ^35^ was used to trim off the adapter sequences and reads were trimmed or filtered according to the quality scores with PRINSEQ ^36^. The high-quality reads were mapped onto the mouse reference genome (mm10) using STAR ^37^ and only uniquely-mapped reads were kept for the downstream analysis. The alignments were further processed using SAMtools ^38^ in terms of manipulation and conversion of BAM files. Read summarisation was conducted and read counts for the genes were generated using featureCounts function of Rsubread package ^39^. Differential expression analysis for compared two sample groups and the likelihood ratio test (LRT) for time-series experiments were done using DESeq2 package ^40^. degPatterns function from DEGreport package was chosen to characterise the patterns of gene expression and cluster genes based on gene expression profiles for 5478 genes (for hard samples) and 8042 genes (for soft samples) that were differentially expressed across time points based on p_adj_ < 0.001 from Likelihood ratio test (LRT) (https://www.bioconductor.org/packages/release/bioc/html/DEGreport.html). DSeq2 performs the statistical analysis between the two groups of samples and provides adjusted p-values (p_adj_) used in all the comparisons here to assess if differences are significantly significant. WebGestalt (WEB-based Gene SeT AnaLysis Toolkit ^41^) was used for gene set overrepresentation analyses under the significance level of FDR < 0.05. Values of RPKM (Reads Per Kilobase of transcript per Million mapped reads) were used to compare expression levels for specific genes of interest. Final graphs were generated using Prism (GraphPad). RPKM values are shown as mean ± S.D.

### Immunostaining and Fusion Index calculation

To compare fusion into myotubes, C1F cells were cultured on hard (glass) and soft (PDMS) cell culture surfaces, prepared as described above, except that glass coverslips were used instead of petri dishes. For hard surfaces, 50µl of 0.01% gelatin in sterile water was added to 13mm diameter round glass coverslips and allowed to set at 37°C. Excess gelatin was aspirated and the glass surfaces were coated with 0.1% collagen solution. For soft surfaces, coverslips were coated with 50µl prepared PDMS as described above, spread carefully with a sterile metal spatula, left to set for 48hrs, treated with sulfo-SANPAH and then coated with 0.1% collagen solution.

C1F myoblasts at passage 3 were seeded onto prepared coverslips at a density of 2×10^5^ cells/ml. For analysis, cells were fixed using 4% paraformaldehyde in phosphate-buffered saline (PBS) at UD, D1, D3 and D7, washed 3x with PBS, and then permeabilised with 0.1% Triton X-100 diluted in PBS containing 1% bovine serum albumin (BSA) (Thermo Fisher Scientific), for 5 minutes. The cells were stained using a primary antibody (A4.1025: which recognises all skeletal muscle myosin heavy chain isoforms ^42^). The antibody was diluted 1 in 10 in wash buffer (PBS containing 1% BSA) together with DAPI (4’6-diaminidino-2-phenylindole) to stain DNA (1/500 dilution), applied to coverslips for 1hr, then removed and coverslips washed 5 times with wash buffer. The secondary antibody (Alexa-Fluor 488 anti-mouse (Thermofisher), diluted 1 in 400, was applied to the coverslips for a further 1hr. The secondary antibody was then removed, coverslips were washed 5 times with wash buffer and once with PBS, then mounted onto glass slides with Prolong® Gold anti-fade mountant (Invitrogen). Slides were left to dry at room temperature in the dark overnight and stored at 4°C before imaging.

To estimate the fusion index, images of the cells were captured using a Delta Vision Widefield Deconvolution Microscope (Delta Vision, USA) using a 40x objective lens (N.A. 1.4). Three biological repeats were performed, and cell images were taken from 10 random fields of view for each repeat. The fusion index was determined by dividing the total number of nuclei found within myotubes (identified by their positive staining for skeletal myosin by the total number of nuclei in the field of view. Values were plotted and compared using Prizm (Graphpad) using multiple unpaired t-tests. Cells were additionally imaged using an LSM880 confocal microscope (Zeiss) and 40x objective lens (N.A 1.4) to image myoblast differentiation in more detail.

To investigate expression of transcription factors, cultures at each time point (UD, D1, D5, D7) were fixed and stained for Pax3 (Developmental studies hybridoma bank (DSHB): diluted 1:20), Pax7 (DSHB, diluted 1:20), MyoD (diluted 1:100, Invitrogen, MA1-41017) and myogenin (diluted 1:50, Invitrogen: MA511486) and co-stained using DAPI, using a similar procedure. To quantify expression, three biological repeats were performed for each time point, on each type of surface, and cell images were taken from 10 random fields of view for each repeat. Numbers of nuclei positively stained for each transcription factor, in each field of view were counted. Values were plotted and compared using Prizm (Graphpad) using multiple unpaired t-tests.

## Acknowledgements

This work was funded by the BBSRC (BB/P007791/1 to M.P. and C.A.J.), BB/M011151/1 Doctoral Training Program (L.R) and MRC grant MR/M000532/1 (to C.A.J.) We acknowledge technical support and advice from Dr Chris Watson, Translational Genomics Unit, Leeds Teaching Hospitals NHS Trust, support from the BioImaging Facility in the Faculty of Biological Sciences, and support from the Wellcome Trust for the Zeiss Airyscan Confocal used in some of the imaging work presented (WT104918MA). RH was funded through an MRC PHD 1233630

## Author contributions

LR acquired and analysed data, DW analysed and interpreted data, RH acquired data, CAJ designed the work, MP conceived and designed the work, analysed and interpreted data. All authors read and commented on the paper.

## Additional information (competing interests)

The authors declare that that they do not have competing financial interests.

## Data availability

RNASeq datasets have been deposited on Sequence Read Archive (SRA) with accession number: PRJNA682314

