## Supplementary Figure S1 for "RNASeq analysis of a Pax3-expressing myoblast clone *in-vitro* and effect of culture surface stiffness on differentiation"

Supplemental Fig. S1

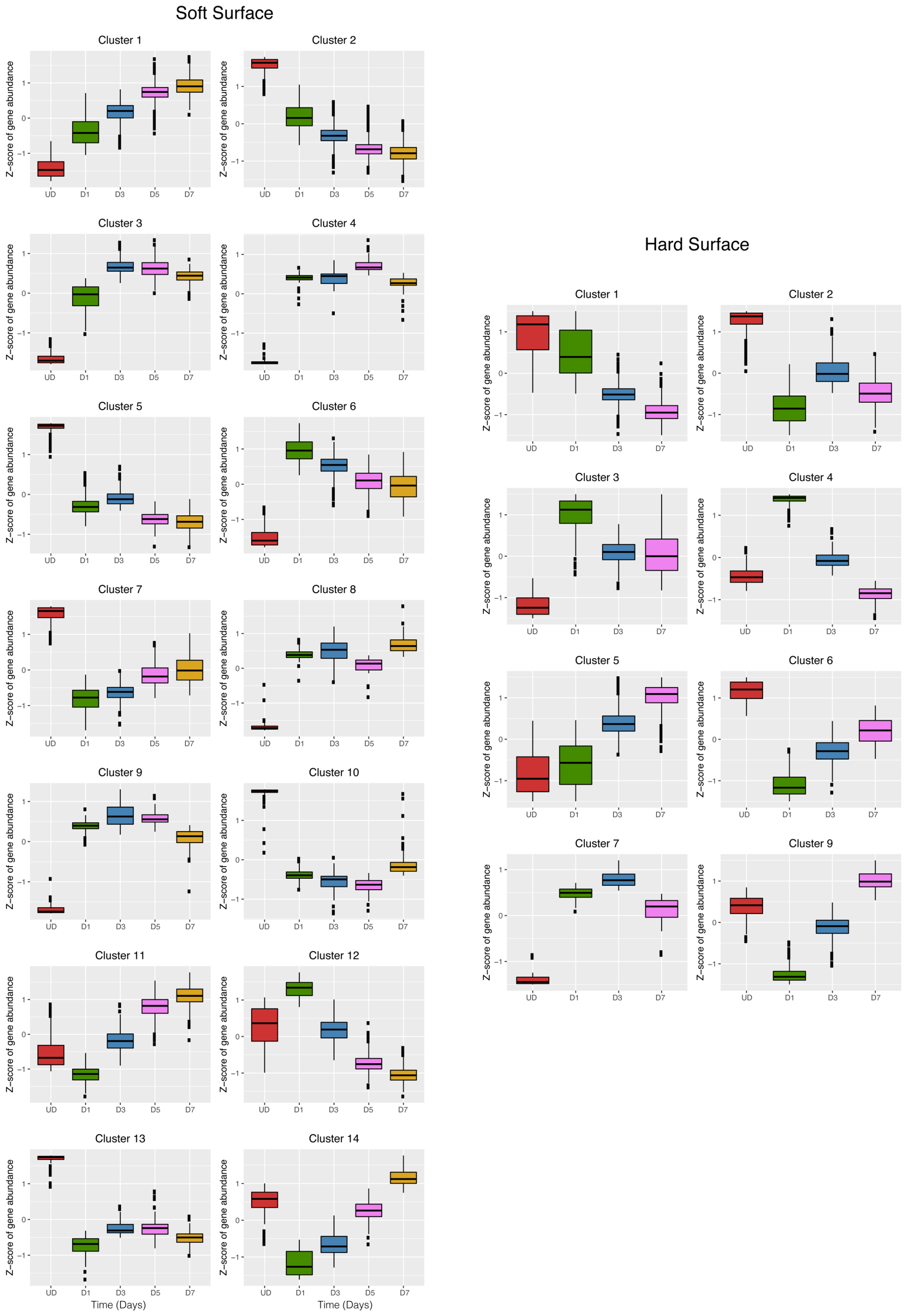


Supplemental Figure S1: Pattern analysis of RNA expression in cells on soft and hard surfaces. The analysis was performed using DegPatterns (see methods).
